# Mouse precision-cut liver and kidney slices: an optimized *ex vivo* model for acute toxicity testing

**DOI:** 10.1101/2025.07.21.665916

**Authors:** Christiane Spruck, Ghizlan Bousejra, Ahmed Erbay, Sonja Herbrich, Andreas Dimitriadis, Zoe Röntgen, Frauke Roolfs, Isaac Musong Mboni-Johnston, Gerhard Fritz, Seddik Hammad, Nicole Schupp

## Abstract

As excretion organs, the kidneys and liver are exposed to high concentrations of potentially toxic substances. While animal models remain the gold standard for organ-specific toxicity testing, alternative *ex vivo* approaches are essential to align with the 3R principles (refinement, reduction, replacement). Precision-cut tissue slices (PCTS) retain native tissue architecture, cellular heterogeneity, the interplay of different cell types, and metabolic capacity, offering a promising link between *in vitro* and *in vivo* models. Here, we aimed to establish an optimized protocol for preparing and culturing precision-cut kidney and liver slices (PCKS and PCLS) from mice for use in substance-oriented toxicological tests. Key parameters - including slice thickness, media composition, oxygenation, glucose levels, and incubation time - were refined to maintain tissue viability and metabolic function. Five known toxins - acetaminophen, cyclosporin A, cisplatin, arsenic trioxide, and aristolochic acid I - were tested. While PCKS showed comparable sensitivity to established kidney cell lines, PCLS achieved IC_50_ values closely matching *in vivo* toxicity data. High reproducibility across different experimenters was achieved, highlighting the robustness of the model. In conclusion, this *ex vivo* system provides a valuable, reproducible, and ethically approved platform for acute nephrotoxicity and hepatotoxicity testing, supporting preclinical drug screening and potentially reducing reliance on animal experiments.

## Introduction

Toxic effects on the kidneys and liver can currently only be determined through animal studies, as there are no recognized alternative methods to fully replace *in vivo* testing. The kidney’s highly complex morphology is difficult to replicate outside the organism. There are various research approaches to develop high-throughput systems using primary cells, cell lines, organoids, or microfluidic chips, however, no breakthrough has yet been achieved [1]. Most advanced systems focus solely on proximal tubular cells - a frequent target of toxicity due to their high transport capacity [2]. However, maintaining this transport capacity *in vitro* is another problem for potential replacement methods. The liver, though structurally less complex than the kidney, also poses difficulties for alternative methods, as current *in vitro* models suffer from poor predictive power due to metabolic deficiencies in available cell lines [3].

In order to comply with the 3R principles (refinement, reduction, replacement), researchers are developing *in vitro* and *in silico* approaches to assess kidney and liver toxicity. [4, 5]. However, the pathophysiology of tissue damage - whether caused by endogenous toxins, environmental pollutants, or drugs - is highly complex, involving multiple cell types and interactions. Reliable risk assessment of organ-specific toxicity requires test systems that closely mimic *in vivo* conditions, including native tissue architecture. This complexity explains why animal models remain the gold standard for addressing fundamental research questions and minimizing regulatory uncertainty. Here, precision-cut tissue slices (PCTS) offer a promising link between *in vitro* and *in vivo* models [6]. Unlike conventional monotypic cell cultures, PCTS retain native tissue architecture, cellular heterogeneity, and the interplay of different cell types, making them particularly valuable for organ-specific toxicity studies. PCTS can be prepared from a variety of organs, including the liver [7], kidney [8], lung [9, 10], heart [11], and intestine [12, 13]. Compared to animal experiments, PCTS enable simultaneous testing of multiple substances in a single organ [14] and the investigation of time- and dose-dependent toxicological effects without the need for large numbers of animals and without subjecting the animals to the treatment. Moreover, the metabolism of foreign substances, which is often rapidly lost in cell cultures, is preserved for several days in PCTS [6].

In this study, existing protocols for the cultivation of PCTS were simplified and improved with the aim of reliably maintaining the viability and functionality of the slices for testing acute toxicity. In addition, the reproducibility of the optimized method was verified. This serves to prepare PCTS toxicity testing for future development as a regulatory-accepted method. Achieving these objectives would advance the 3Rs by reducing reliance on animal studies while providing more physiologically relevant toxicity data for drug development and environmental risk assessment.

## Materials and Methods

If not stated otherwise, all chemicals and media were either from Merck, St. Louis, MO, USA, or ThermoFisher Waltham, MA, USA.

### Animals

The tissue sections used for toxicity testing and immunohistochemical analyses were obtained from male C57BL/6J wild-type mice aged 10-16 weeks, provided either by Janvier Laboratories (Le Genest-Saint-Isle, France) as well as from the internal breeding of the Central Institution for Animal Research and Scientific Animal Welfare (ZETT) of Heinrich Heine University (Düsseldorf, Germany). All animals used for organ collection were registered under the number O55/16 of the ZETT of the Heinrich Heine University (Düsseldorf, Germany). Following deep anaesthesia with ketamine (120 mg/kg, Zoetis, Berlin, Germany) and xylazine (8 mg/kg, i.m., Elanco Deutschland GmbH, Bad Homburg, Germany), the animals were euthanized by cervical dislocation. Both kidneys and the liver were immediately harvested in strict accordance with the European Community guidelines for the care and use of laboratory animals and the German Animal Welfare Act. The organs were stored in ice-cold Belzer UW® solution (Bridge to Life™ Ltd., Northbrook, IL, USA) until further processing for tissue slicing.

### Cell lines

NRK-52E cells (CRL-6509, ATCC, Manassas, VA, USA), normal proximal tubular epithelial cells isolated from Rattus norvegicus [15], were cultured in DMEM high glucose, supplemented with 10% FCS and 1% Penicillin-Streptomycin at 37 °C, 21 % O_2_ and 5 % CO_2_. HepG2 cells (HB-8065, ATCC, Manassas, VA, USA) derived from liver tumor tissue of a 15-year-old male patient [16], were cultured in RPMI, supplemented with 10% FCS and 1% PenStrep under the same conditions. Proximal tubular-like cells (PTELC) differentiated from induced pluripotent stem cells (iPSC) were generated and maintained as described by Mboni-Johnston et al. [17].

### Preparation of precision cut tissue slices

Ice-cold Krebs-Henseleit-Buffer (KHB [18], supplemented with 2.5 g glucose/l and 10 mM HEPES) was used in the Alabama R&D Tissue Slicer (Alabama Specialty Products, Inc., Munford, AL, USA) for tissue slicing. Kidneys were embedded in agarose, placed in the organ holder and subsequently sliced. For liver preparation, 8 mm biopsy punches were taken and then sliced. Precision-cut kidney slices (PCKS) and precision-cut liver slices (PCLS) of comparable size (small kidney slices comprising only the cortex were discarded) were washed in KHB and stored in cold Belzer UW® solution on ice until further use.

### Cultivation of precision cut tissue slices

PCKS and PCLS were cultivated in 12-well plates at 37 °C, 21 % O_2_ and 5 % CO_2_ on a shaker to enhance oxygen availability. In the initial experiments, PCKS were incubated in Williams E Medium, supplemented with 2 mM Glutamax, 15 mM HEPES, 2.5 g glucose/l (resulting in a total of 4.5 g glucose/l) and 10 mg ciprofloxacin/l. Later in the project, the culture medium was optimized to improve tissue viability. Slices were incubated in DMEM without glucose, either with (for PCLS) or without (for PCKS) phenol red, supplemented with 2 mM Glutamax, 2 g/l glucose, 25 mM HEPES, 2x Insulin-Transferrin-Selenium Supplement, 5% FCS, to improve viability. The optimal slice thickness after the evaluation was 200 µm for PCKS and 250 µm for PCLS.

### Viability assays

Viability of cultured cells and of cells in PCTS was measured with the MTT assay, which is based on the reduction of water-soluble, yellow 3-(4,5-dimethylthiazol-2-yl)-2,5-diphenyltetrazolium bromide (MTT) to insoluble, purple formazan by cellular reductases. This can be quantified colorimetrically after dissolving the formazan in an organic solvent [19]. Cells were seeded in quadruplicates in 96 well plates. After 24 h exposure to the various toxins, 20 µl MTT solution (5 mg/ml) was added to the medium and incubated for about 40 min at 37 °C. The supernatant was removed and the blue formazan crystals were dissolved in 100 µl dimethyl sulfoxide (DMSO) for 5 min at room temperature on a shaker. For tissue slices, 260 μl of the MTT solution (5 mg/ml) was added to the culture medium and incubated for 1 hour at 37 °C and 5 % CO₂ on a shaker. After incubation, the tissue slices were carefully weighed and transferred to 1 ml of isopropanol to dissolve the formazan by shaking at room temperature for 30 minutes. Subsequently, 100 μl of the isopropanol-formazan solution was pipetted in quadruplicate into a 96-well plate and absorbance was measured at 560 nm using a spectrophotometer. The absorbance values were normalized to the wet weight of the tissue slices and technical replicates were averaged. The viability of untreated cells and PCTS was defined as 100%, allowing viability under different conditions to be compared. Viability of PTELC was assessed with the Alamar Blue assay, in which non-fluorescent resazurin is reduced to fluorescent resorufin by aerobic respiration of metabolically active cells, as previously described [17].

As a second cytotoxicity assay to evaluate viability of PCTS, the ATP assay, which is based on the conversion of luciferin to bioluminescent oxyluciferin by the enzyme luciferase in the presence of ATP produced by functional mitochondria [20], was used. The resulting luminescence is directly proportional to the ATP content and thus reflects the viability and functionality of the tissue [21]. Tissue slices were weighed after treatment and stored at -80 °C until analysis. Following thawing, tissue samples were lysed using the Tissue Lyser II (Qiagen, Hilden, Germany). ATP levels were quantified in the supernatant using the ATPlite Luminescence Assay System Kit (Perkin Elmer, Waltham, MA, USA), according to the manufacturer’s instructions. Luminescence values were normalized to the wet weight of the slices, and the ATP level of the untreated control was defined as 100% viability.

As a second cytotoxicity assay for the cell lines, the neutral red assay was conducted, which is based on the principle of an ion trap, where the dye neutral red is trapped in intact lysosomes in the cells. The cells were seeded in quadruplicates in 96 well plates and after the 24 h treatment, the medium was replaced with 200 µl neutral red solution (10 % neutral red stock (0.1 % (w/v) neutral red in distilled water, 10 % HEPES, 80 % medium). After 1 h incubation at 37 °C, the cells were fixed by replacing the neutral red solution with the fixing solution (1 % formaldehyde, 1 % CaCl_2_ in distilled water), washed with 200 µl phosphate-buffered saline and the dye was extracted with the extraction solution (50 % ethanol, 1 % acetic acid in distilled water) for 5 min at room temperature on a shaker. Absorption was measured at 540 nm using a spectrophotometer. Absorption values of the untreated cells were defined as 100% viability.

### Histological staining

For the hematoxylin eosin (HE) staining, paraffin embedded kidney and liver sections of 3 μm thickness were prepared, mounted on glass slides and incubated for 60 min at 60 °C. The sections were deparaffinized, stained for 5-30 s with hematoxylin, rinsed for 8 min under flowing tap water to make the cell nuclei visible and stained with eosin (Carl Roth GmbH + Co. KG, Karlsruhe, Germany) for 2-3 min. After dehydration, the sections were mounted with Entellan, and were analyzed microscopically (RM 2164, Leica, Wetzlar, Germany) at a 200 x magnification.

### Statistical Analysis

Statistical analysis was performed using GraphPad Prism version 6 (GraphPad Software, San Diego, CA, USA). Data are presented as mean ± standard deviation (SD) of at least three independent experiments (n ≥ 3). Normal distribution was checked using the Kolmogorov-Smirnov test. For comparisons of two normally distributed groups the two-tailed unpaired Student’s t-test was employed. For non-normally distributed values, the Mann-Whitney U-test was used. Multiple groups were tested by one-way analysis of variance (ANOVA) with Tukey’s or Dunnett’s post hoc test. Non-parametrical datasets of multiple groups were analyzed with Kruskal-Wallis one-way ANOVA on ranks test. Statistically significant differences between the groups were assumed at a p-value ≤0.05.

## Results

### Optimization of slice thickness, medium composition, and culture duration

To establish optimal culture conditions for murine PCKS and PCLS, we first evaluated slice thickness. As shown in Figure 1, the viability of the PCKS was strongly dependent on slice thickness. Slices up to 200 µm thickness retained viability after 24 hours; however, a significant reduction was observed at a thickness of 300 µm. After 48 hours, slices with a thickness of 200 µm also showed markedly decreased viability. In contrast, PCLS viability was less affected by slice thickness, with only the 250 µm thick slices showing a significant reduction in metabolic activity at 48 hours compared to the control at 0 hours.

**Figure 1:**
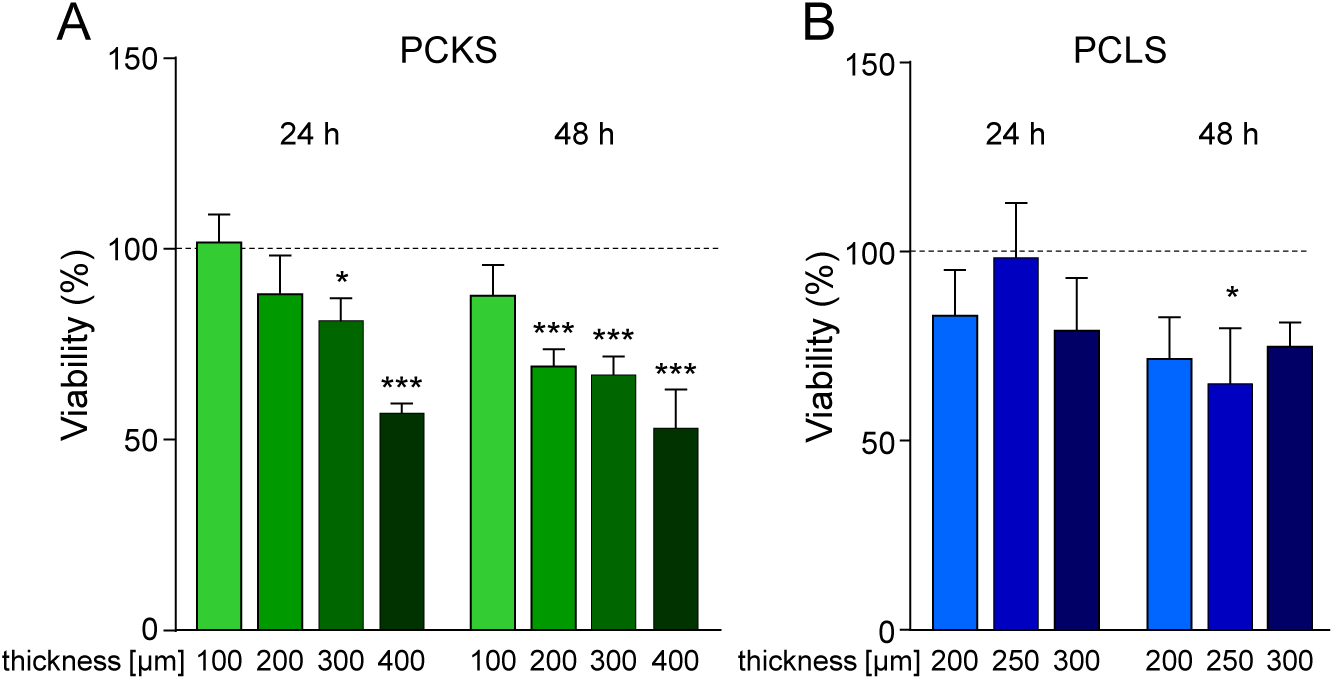
Influence of slice thickness on viability of precision-cut tissue slices. Kidneys (A) and livers (B) of the mice were cut into slices of the specified thickness, and their viability was determined after 24 and 48 h by measuring metabolic activity in the MTT assay. Data are presented as mean + SEM (n = 3, N = 3, with n = independent and N = technical replicates). *p≤0.05, ***p<0.001 vs. slices of the respective thickness at 0 h (one-way ANOVA). MTT = 3-(4,5-dimethylthiazol-2-yl)-2,5-diphenyltetrazolium bromide, PCKS = precision cut kidney slices, PCLS = precision cut liver slices.

To further improve the viability and functionality, we next investigated cultivation conditions for PCTS described in previous studies. Enriching the slicing buffer and the cultivation medium with 95% oxygen did not improve viability in our study. In PCKS, a significant temporary increase in metabolic activity was observed after 4 hours (Figure 2A), while PCLS showed a tendency toward reduced viability under hyperoxic conditions (Figure 2B).

**Figure 2:**
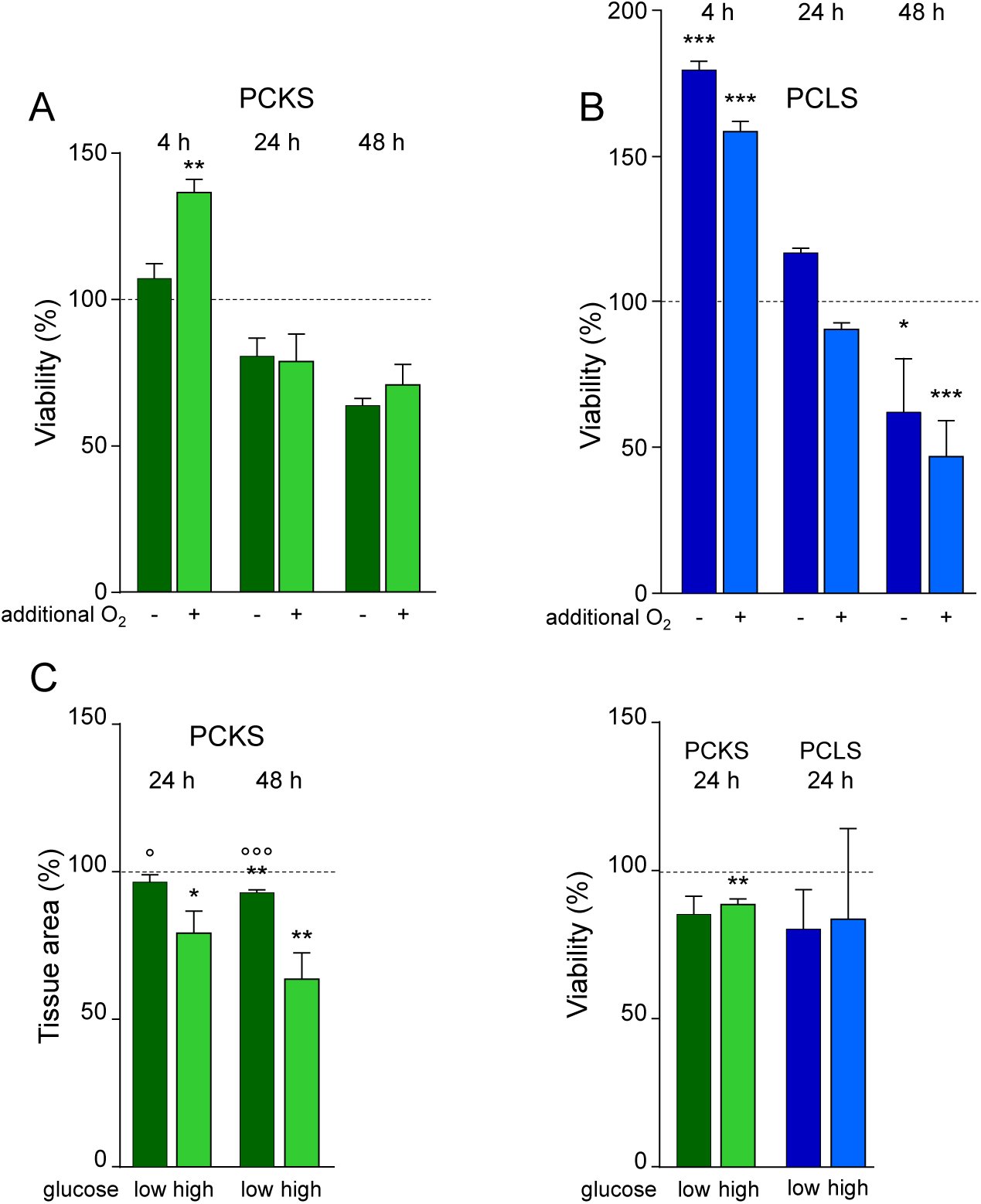
Effect of oxygenation and glucose concentration on PCTS viability. Kidneys (A) and livers (B) of mice were either cut in non-oxygenized (-) or oxygenized (+) organ storage buffer and incubated in non-oxygenized (-) or oxygenized (+) medium for up to 48 h. Afterwards, the viability of the slices was determined by measuring the metabolic activity in the MTT assay. (C) Surface area of PCKS was measured after incubation for 24 and 48 h in high (4.5 glucose/l) or low (2 g glucose/l) glucose media. (D) Viability after 24 h incubation in high and low glucose medium was determined by MTT assay. Data are presented as mean + SEM (n = 3, N = 3). *p≤0.05, **p<0.01, ***p<0.001 vs. 0 h (two-way ANOVA), ° p≤0.05, °°°p<0.001 vs. high glucose (one-way ANOVA). MTT = 3-(4,5-dimethylthiazol-2-yl)-2,5-diphenyltetrazolium bromide, PCKS = precision cut kidney slices, PCLS = precision cut liver slices, PCTS = precision cut tissue slices.

A closer examination of PCKS revealed that they had shrunk significantly after 24 hours of incubation in the initial high glucose medium (Figure 2C). Reducing glucose levels to physiological values mitigated the shrinkage of the tissue sections, so that it only became significant after 48 hours in the optimized medium (Figure 2C). Notably, viability of PCKS and PCLS in low-glucose conditions was comparable to that under high-glucose conditions (Figure 2D), supporting the use of physiological glucose concentration in subsequent experiments.

Under optimized conditions (non-oxygenated media and physiological glucose), viability loss over time was evaluated. PCKS retained approximately 70% viability at 24 hours, 50% at 48 hours, and 25% at 96 hours (Figure 3A). In contrast, PCLS, maintained approximately 75 % viability throughout 96 hours (Figure 3B).

**Figure 3:**
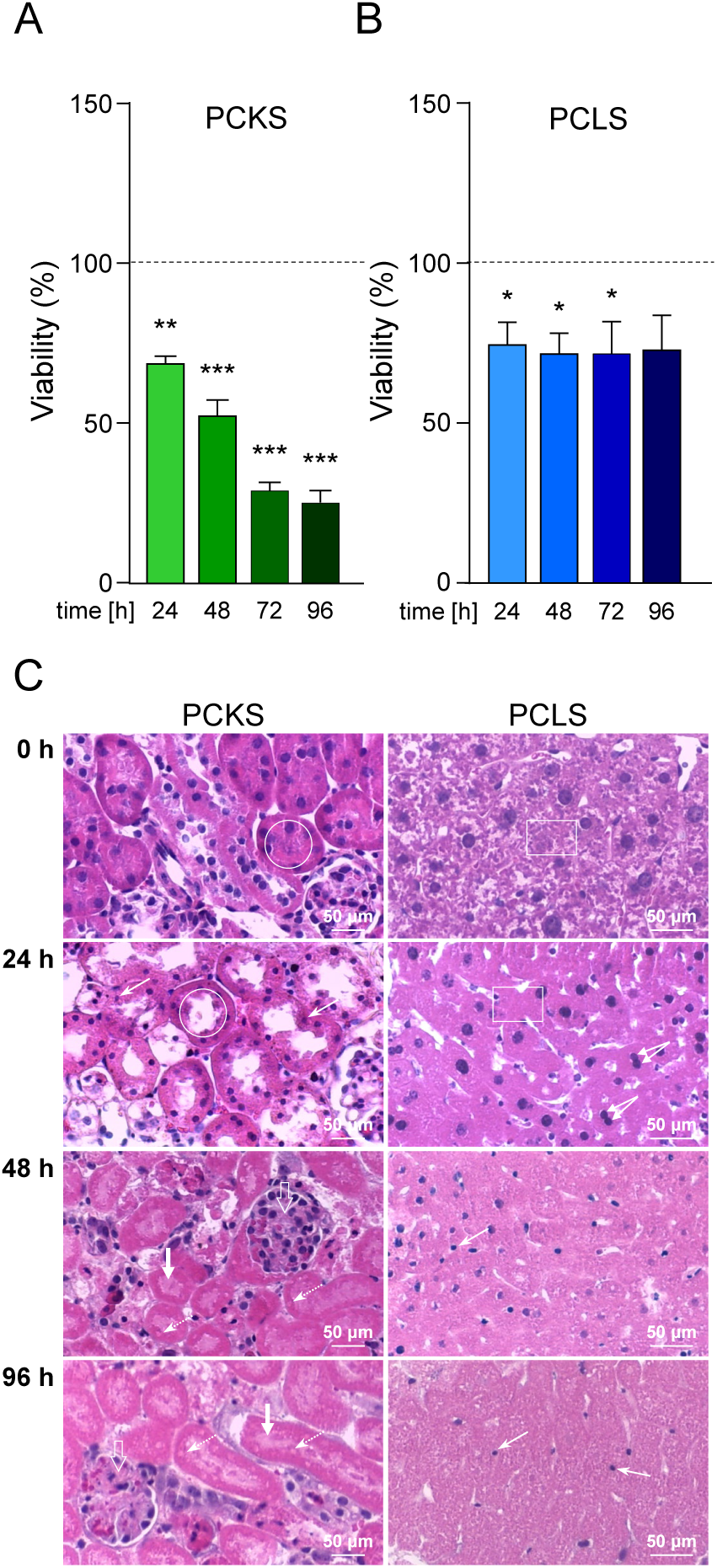
Viability of PCTS over an incubation period of 96 hours. Viability of PCKS (A) and PCLS (B) under optimized incubation conditions was measured using the MTT assay. (C) Representative images of HE-stained PCKS and PCLS. White circle 0 h: brush border of proximal tubule, white circle 24 h: lost brush border of proximal tubule, small arrow: condensed nuclei, small dashed arrow: outline of a nuclei, big open arrow: glomerular swelling, big filled arrow: luminal obstruction, white rectangle 0 h: granular liver structure, white rectangle 24 h: loss of granularity, double arrow: binucleated hepatocyte. Data are shown as mean + SEM (n = 3, N = 3). *p≤0.05, **p<0.01, ***p<0.001 vs. 0 h (one-way ANOVA). HE = hematoxylin and eosin, MTT = 3-(4,5-dimethylthiazol-2-yl)-2,5-diphenyltetrazolium bromide, PCKS = precision cut kidney slices, PCLS = precision cut liver slices, PCTS = precision cut tissue slices.

Histological analysis via HE-staining (Figure 3C) confirmed the results of the viability assay. PCKS showed intact morphology immediately after slicing. After 24 hours, tubular cell nuclei were condensed and brush borders of the proximal tubule were lost. After 48 and 96 hours, glomerular swelling and luminal obstruction were evident. Furthermore, no cell nuclei were stained in the tubules, but since the outlines of the nuclei are still clearly visible, this is most likely due to a technical problem during staining. PCLS exhibited characteristic lobular architecture and granular cytoplasm at 0 hours (Figure 3C). By 24 hours, cytoplasmic granularity was diminished and nuclear compaction was apparent. Binuclear hepatocytes – indicative of regenerative activity – were observed at the same time. At 48 and 96 hours onwards, progressive nuclear condensation and loss of nuclei were noted.

Given the restricted viability of PCKS and since the substances used are all able to cause acute damage to tissues, the subsequent toxicity experiments were limited to 24-hour exposures.

### Organotypic toxicity profiling of five reference toxins in PCKS and PCLS compared to renal and hepatic cell lines

PCKS and PCLS were treated with five toxins known to affect renal and/or hepatic function. Comparisons were made to cellular models - NRK (normal rat kidney cells) and HepG2 (human hepatoma cells). Two viability tests were applied to both tissue slices (MTT and ATP assay) and cell lines (MTT and neutral red uptake (NR) assay). Acetaminophen exposure induced a comparable, concentration-dependent viability loss in PCKS and NRK cells, with similar IC_50_ values, except for a higher IC_50_ in the NR assay for NRK cells (Figure 4A). PCLS showed heightened sensitivity to acetaminophen, with IC_50_ values around 1 mM, while HepG2 cells only displayed moderate sensitivity (Figure 4B). Upon treatment with cyclosporin A, all models exhibited similar IC_50_ values, except for PCKS, PCLS and HepG2 assessed via the MTT assay, where the IC_50_ was notably higher (Figures 4 C and D). In addition, proximal tubular cells (PTELC) differentiated from hiPSC showed similar sensitivity as NRK cells. Exposure to cisplatin, resulted in an assay-dependent variability. ATP- and NR-based assessments indicated greater sensitivity compared to MTT-derived values (Figures 4E and F). PTELC, HepG2 cells (NR assay), and PCLS (ATP assay) demonstrated the highest sensitivity. PCKS showed an ATP assay-based IC_50_ approximately twice that of PCLS. For arsenic trioxide, ATP assay results indicated high sensitivity in both PCKS and PCLS, with IC_50_ values between 4–8 µM (Figures 4G and H). However, MTT-derived IC_50_ values were 4–6-fold higher, suggesting reduced sensitivity of the MTT assay. In contrast, HepG2 cells exhibited much lower sensitivity across both viability assays. Aristolochic acid I induced pronounced cytotoxicity only in PCLS (Figures 4I and J). In conclusion, treatment with five reference toxins revealed that PCKS and PCLS generally exhibited toxicity profiles consistent with their respective cellular models, with assay-dependent differences in sensitivity; notably, PCLS showed heightened sensitivity to acetaminophen, cisplatin and aristolochic acid, while PCKS and kidney-derived cells demonstrated strong responses to cyclosporin A and arsenic trioxide, highlighting the value of tissue slices for organ-specific toxicity assessment.

**Figure 4:**
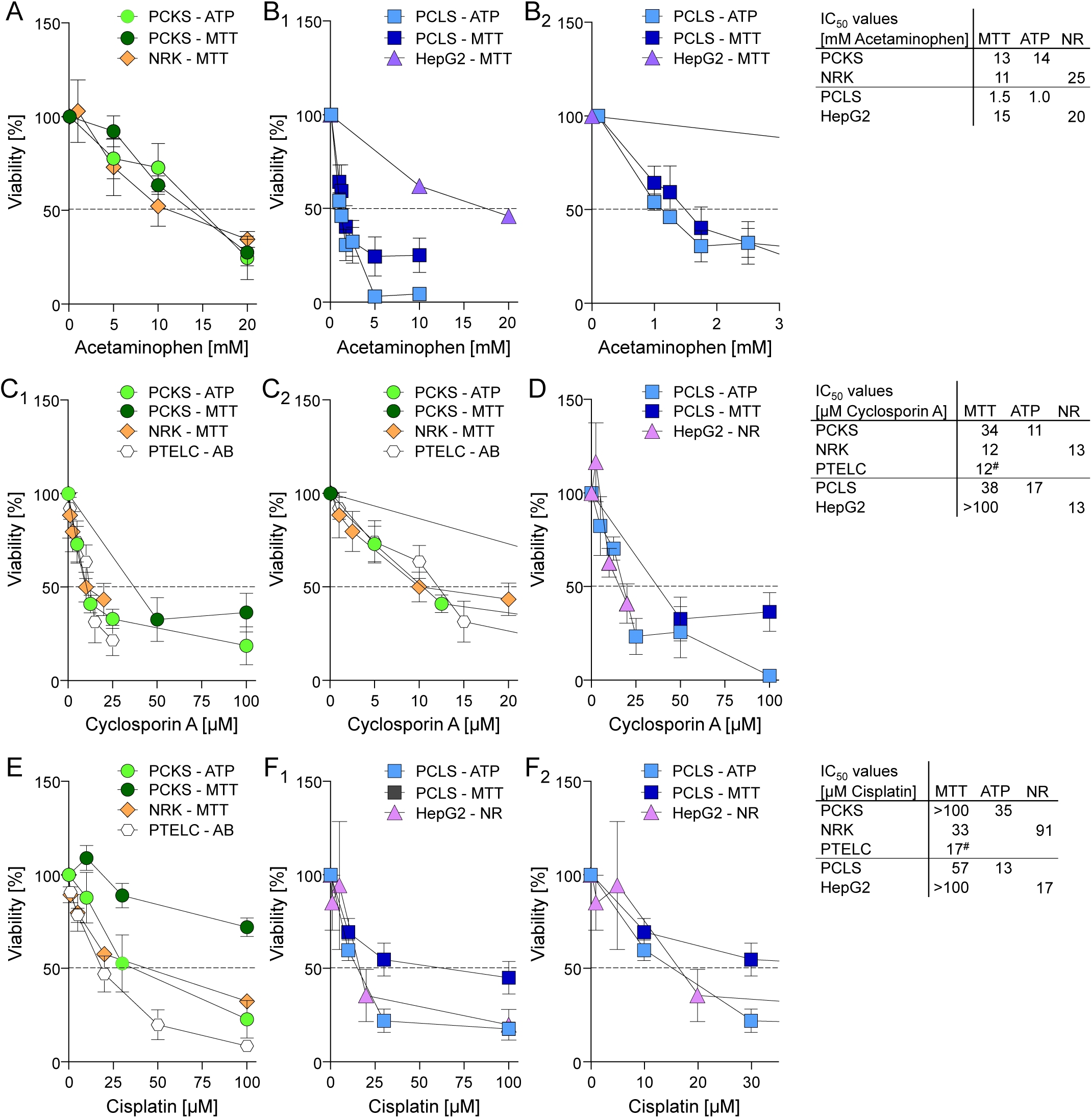
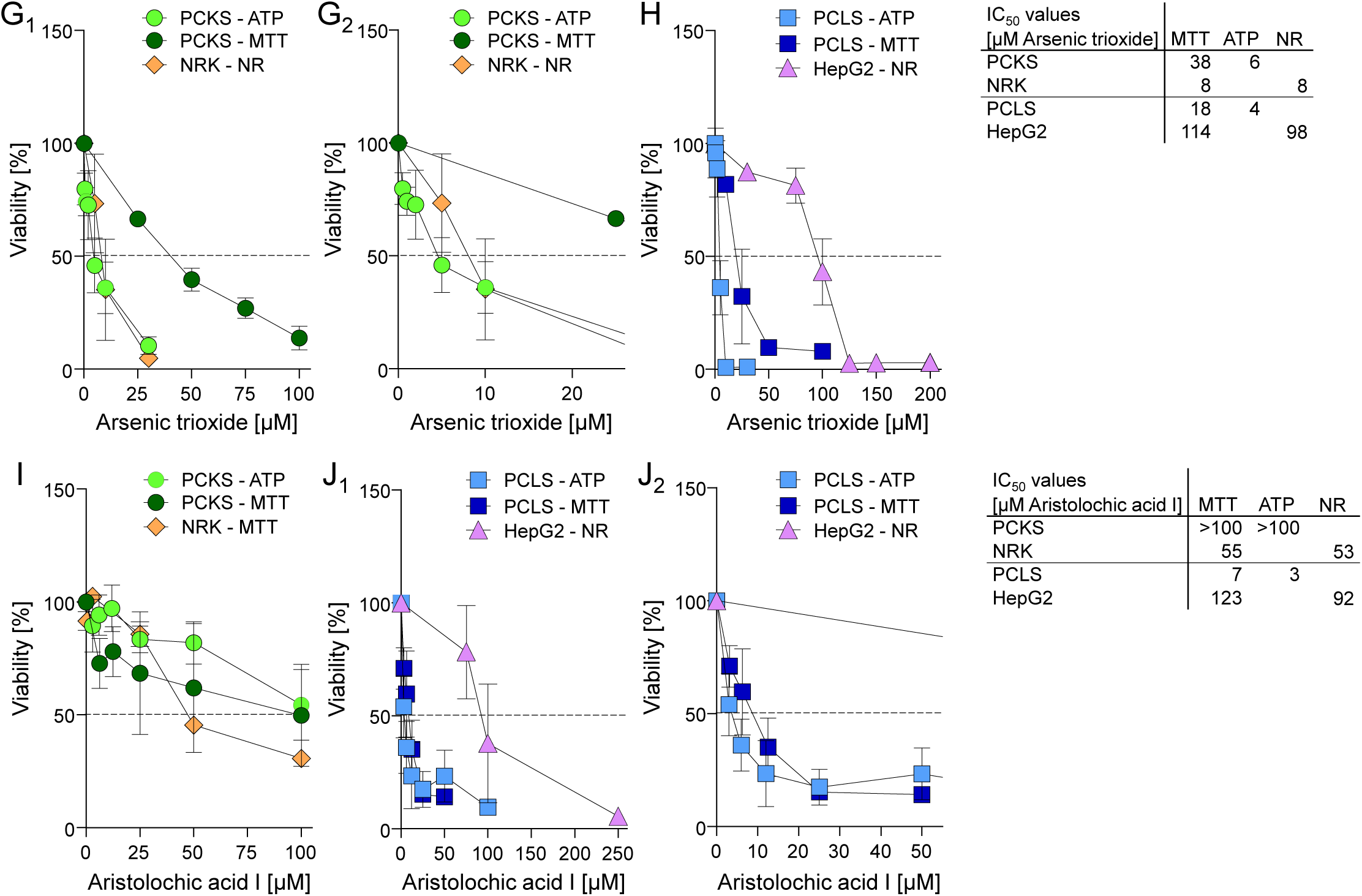
Viability of PCKS and PCLS as well as kidney and liver cells after treatment with drugs and toxins. PCKS and NRK cells (A) and PCLS and HepG2 cells (B_1_ and B_2_ = section of the graph at low concentrations) were treated for 24 h with acetaminophen. Viability was measured either with the ATP or the MTT assay. Additionally, the calculated IC_50_ values of the different cell and tissue models are shown in a table. PCKS and NRK cells (C_1_ and C_2_ = section of the graph at low concentrations) and PCLS and HepG2cells (D) were treated for 24 h with cyclosporin A. Viability was measured using the ATP, the MTT, the NR or the ^#^Alamar Blue assay. Additionally, the calculated IC_50_ values of the different cell and tissue models is shown in a table. PCKS and NRK cells (E) and PCLS and HepG2cells (F_1_ and F_2_ = section of the graph at low concentrations) were treated for 24 h with cisplatin. Viability was measured using the ATP, the MTT, the NR or the ^#^Alamar Blue assay. Additionally, the calculated IC_50_ values of the different cell and tissue models are shown in a table. The dotted line indicates 50 % viability, and therefore, the IC_50_. Data are given as mean (n = 3-7, N = 3) with SEM when tissue slices were analyzed and SD when cells were analyzed. AB = Alamar Blue assay, ATP = adenosine triphosphate assay, HepG2 = human liver cancer cell line, IC_50_ = half maximal inhibitory concentration, MTT = 3-(4,5-dimethylthiazol-2-yl)-2,5-diphenyltetrazolium bromide assay, NR = neutral red assay, NRK = normal rat kidney cell line, PCKS = precision-cut kidney slices, PCLS = precision-cut liver slices, PTELC = proximal tubular epithelial-like cells. PCKS and NRK cells (G_1_ and G_2_ = section of the graph at low concentrations) and PCLS and HepG2cells (H) were treated for 24 h with arsenic trioxide. Viability was measured using the ATP, the MTT or the NR assay. Additionally, the calculated IC_50_ values of the different cell and tissue models are shown in a table. PCKS and NRK cells (I) and PCLS and HepG2cells (J_1_ and J_2_ = section of the graph at low concentrations) were treated for 24 h with aristolochic acid I. Viability was measured using the ATP, the MTT or the NR assay. Additionally, the calculated IC_50_ values of the different cell and tissue models are shown in a table. The dotted line indicates 50 % viability, and therefore, the IC_50_. Data are given as mean (n = 3-7, N = 3) with SEM when tissue slices were analyzed and SD when cells were analyzed. ATP = adenosine triphosphate assay, HepG2 = human liver cancer cell line, IC_50_ = half maximal inhibitory concentration, MTT = 3-(4,5-dimethylthiazol-2-yl)-2,5-diphenyltetrazolium bromide assay, NR = neutral red assay, NRK = normal rat kidney cell line, PCKS = precision-cut kidney slices, PCLS = precision-cut liver slices, PTELC = proximal tubular epithelial-like cells.

**Figure 5:**
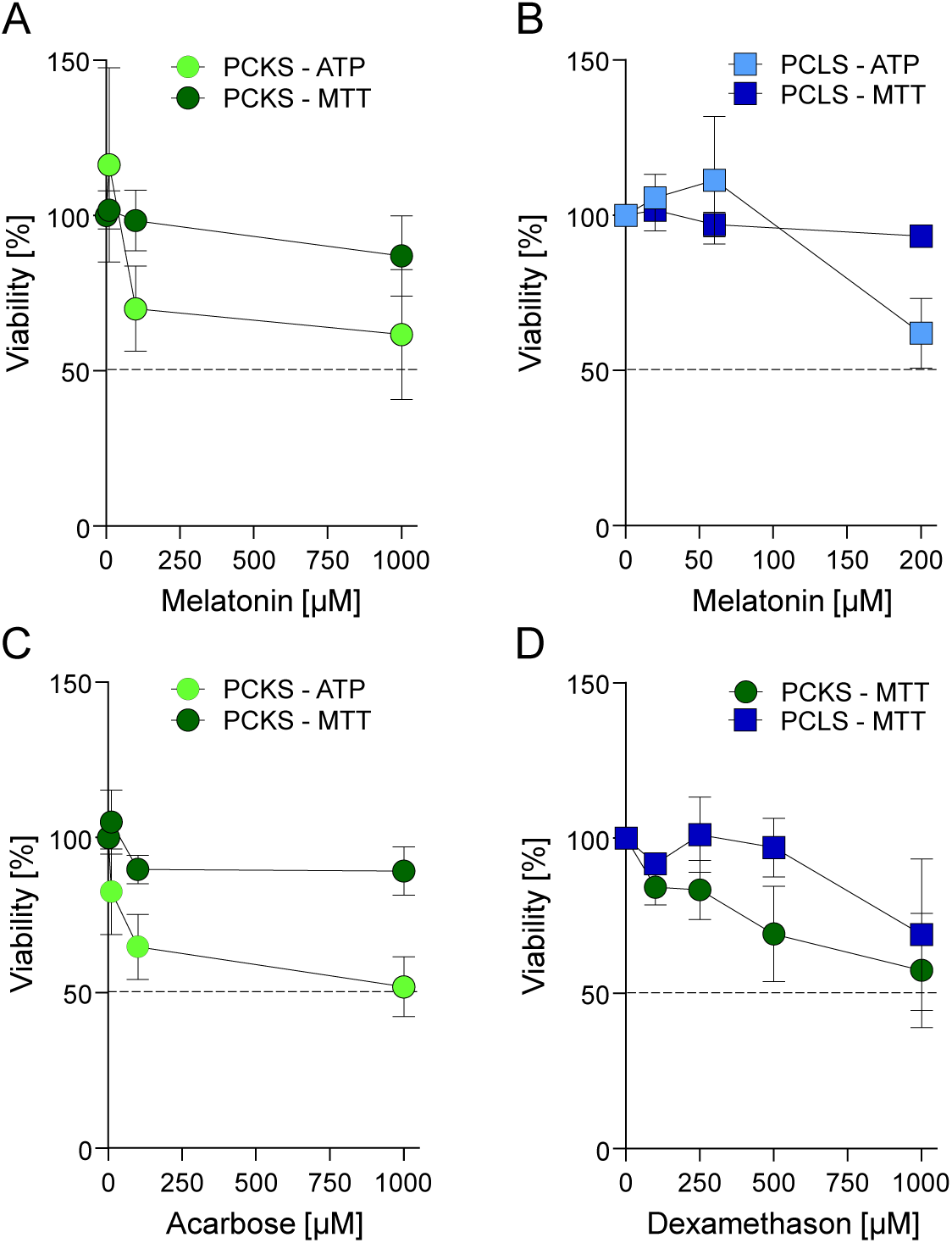
Viability of PCKS and PCLS after treatment with potential negative control substances. PCKS (A) and PCLS (B) were treated with melatonin, and viability was measured with ATP and MTT assay. (C) PCKS were treated with acarbose, and viability was measured with ATP and MTT assay. (D) PCKS and PCLS were treated with dexamethasone, and viability was measured with MTT assay. The dotted line indicates 50 % viability, and therefore, the IC_50_. Data are given as mean ± SEM (n = 3-5, N = 3). ATP = adenosine triphosphate assay, IC_50_ = half maximal inhibitory concentration, MTT = 3-(4,5-dimethylthiazol-2-yl)-2,5-diphenyltetrazolium bromide assay, PCKS = precision-cut kidney slices, PCLS = precision-cut liver slices.

### Evaluation of potential negative control substances

Three substances - melatonin, acarbose (only on PCKS, as acarbose has been linked to rare liver injuries [22]), and dexamethasone - were evaluated as potential negative controls. MTT assay results confirmed minimal toxicity for all three, except dexamethasone, which reduced viability in both PCKS and PCLS. However, ATP assay revealed a reduction in viability for all compounds, with acarbose causing an almost 50% decrease, with the result that none of the substances is suitable as a negative control.

### Demonstration of reproducibility

Acetaminophen served as a positive control in all experiments, allowing us to obtain results for organ slices treated with this substance from multiple experimenters. As shown in Figure 6, the concentration curves are almost identical, except for the early determinations using the MTT assay. This indicates that the results of three different experimenters, even using two different viability assays, proved to be highly reproducible. The three consistent curves yielded an IC_50_ of around 14 mM for the PCKS and an IC_50_ of around 0.9 mM for the PCLS. The two deviating MTT IC_50_ values were approximately twice as high.

**Figure 6:**
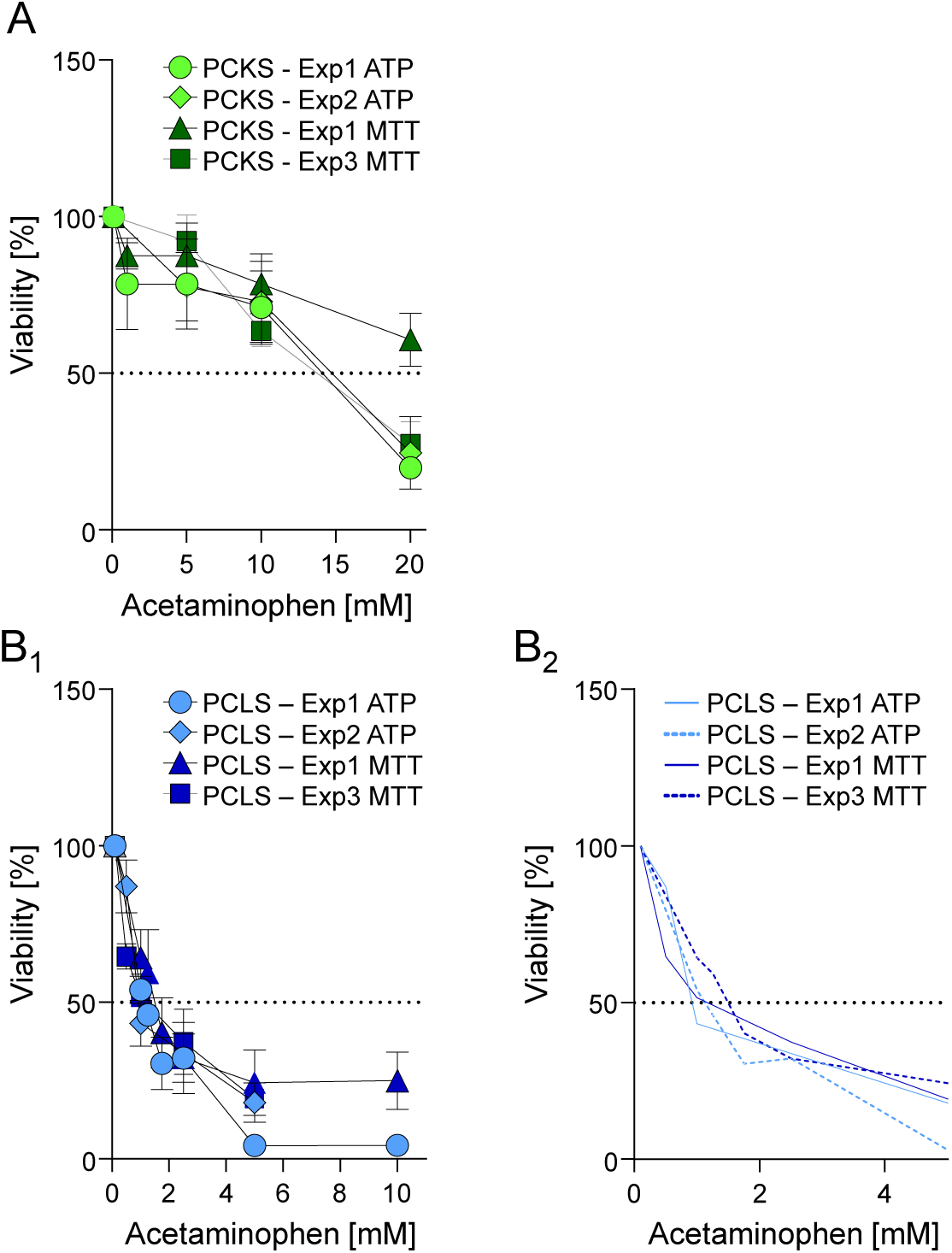
Reproducibility of results between experimenters and viability assays. PCKS (A) and PCLS (B_1_ and B_2_ = section of the graph at low concentrations) were each treated by three different experimenters over the time course of two years with acetaminophen. Viability was measured using either the ATP or the MTT assay. The dotted line indicates 50 % viability and, therefore, the IC_50_. Data are given as mean ± SEM (n = 3-6, N = 3). ATP = adenosine triphosphate assay, IC_50_ = half maximal inhibitory concentration, MTT = 3-(4,5-dimethylthiazol-2-yl)-2,5-diphenyltetrazolium bromide assay, PCKS = precision-cut kidney slices, PCLS = precision-cut liver slices, Exp1-Exp 3 = Experimenter 1-3.

## Discussion

In this study, we aimed to determine whether PCKS and PCLS are suitable models for testing nephro- and hepatotoxicity. To achieve this, we developed an accessible and straightforward protocol for tissue slicing and cultivation that requires only standard laboratory equipment - except the device to prepare the slices - and validated it using a panel of model substances with known toxicological profiles.

Slice thickness is a critical determinant of tissue viability and functional integrity. Consistent with previous reports using PCKS thicknesses ranging from 200 to 310 µm [23-25], we systematically assessed slices between 100 and 400 µm. While 100 µm slices maintained high viability for up to 48 hours, their size was insufficient for further functional analyses. Conversely, 200 µm slices balanced viability and size, with significant viability reduction only after 48 hours. Accordingly, 200 µm was selected for PCKS. For PCLS, where literature frequently reports optimal thicknesses between 200 and 300 µm [26-28], we found no significant viability differences across 200, 250, and 300 µm. Therefore, a thickness of 250 µm - commonly adopted in several studies - was chosen for further PCLS experiments.

A major advantage of the current approach is the elimination of technical barriers related to high oxygenation. While many protocols employ 80-95% oxygen during incubation [23, 28, 29], our results indicate no significant benefit from hyperoxic conditions. On the contrary, supra-physiological oxygen levels risk generating excessive reactive oxygen species (ROS), which may compromise cell physiology [30]. Similarly, the often-used [23, 28, 31, 32] high glucose concentrations (25 mM), corresponding to plasma glucose levels in diabetes [33], failed to confer improved viability. Instead, they induced tissue shrinkage, likely reflecting a stress response analogous to hyperglycemia-induced tissue injury. Consequently, for further experiments, we adopted 11 mM glucose, which is closer to physiological hepatic glucose levels [34], and sufficient to maintain slice viability.

Previous studies primarily relied on ATP content and lactate dehydrogenase (LDH) release for viability assessment [23]. However, variability in reported ATP levels - ranging from stable over 72 hours [23] to paradoxically increased [8, 35, 36], prompted us to include the MTT assay for cross validation. Using our optimized protocol, we achieved prolonged viability with PCLS, retaining approximately 75 % as long as 96 hours, while this value could only be maintained for 24 hours with PCKS. Discrepancies between ATP levels and histological damage shown in other studies [8, 37], underscore the necessity for parallel functional and morphological assessments.

Morphological deterioration was tissue-specific. PCKS showed progressive loss of brush borders in proximal tubules, luminal obstruction with cellular debris, and eventual tubular disintegration - findings consistent with earlier observations [38]. PCLS exhibited cytoplasmic granularity loss and apoptosis-like nuclear condensation within 24 hours. Interestingly, the observed increase in binucleated hepatocytes aligns with regenerative responses documented following liver injury and partial hepatectomy [39], suggesting that the slicing procedure may activate a regenerative program analogous to *in vivo* liver repair. This increase in binucleated cells in PCLS is consistent with other studies showing proliferation in liver slices [37, 40]. Although earlier studies reported the onset of necrosis in both kidney and liver slices [38, 41], the condensed cell nuclei we found indicate predominantly apoptotic cell death.

Both PCKS and PCLS demonstrated compound-specific sensitivities reflective of their respective organ toxicities. Acetaminophen induced a concentration-dependent viability reduction in both tissues, with PCKS (IC₅₀ ∼13 mM) exhibiting lower sensitivity compared to PCLS (IC₅₀ ∼1 mM). PCKS sensitivity was close to that determined in NRK and HepG2 cells and comparable to literature values obtained in human kidney cells (HK-2) and murine PCKS [42, 43]. The IC_50_ value of PCLS aligns with previous reports indicating that mouse PCLS are particularly sensitive to acetaminophen-induced hepatotoxicity [44, 45], potentially due to higher expression of CYP-mediated bioactivation pathways compared to other species [46]. Considering that toxic effects of acetaminophen start at plasma concentrations of 1-2 mM [47-49], mouse PCLS may offer a conservative model for predicting acetaminophen toxicity. The observed toxicity in PCKS further confirms that renal metabolic enzymes remain functional during the early incubation period, consistent with enzyme half-lives ranging from 24 to 140 hours reported in human PCLS [50]. While HepG2 cells were less sensitive to acetaminophen than PCLS, their IC_50_ values were comparable to previous studies [51], underscoring their continued utility despite known limitations in xenobiotic metabolism. In the present study, we used HepG2 cells, recognizing that while they do not replicate the phenotype and function of primary human hepatocytes as closely as HepaRG cells - an emerging and promising alternative for in vitro liver models - they are not consistently outperformed across all toxicological applications [52, 53]. Moreover, the use of HepaRG cells is limited by complex and time-consuming differentiation protocols, as well as variability in reported outcomes [54], making them less suitable for our objective of establishing a broadly applicable platform for PCTS implementation across multiple laboratories.

Cyclosporin A exhibited similar IC_50_ values across PCKS, PCLS, and kidney or liver-derived cell lines, ranging between 10-20 µM. These values are consistent with *in vitro* reports [55-57] and correspond to tissue concentrations observed *in vivo*, which can exceed plasma levels (0.25-0.8 µM) by up to 50-fold [58-60]. This suggests that both PCKS and PCLS faithfully recapitulate cyclosporin A-induced cytotoxicity relevant to clinical exposure levels.

Cisplatin exhibited the highest cytotoxicity in PCLS (measured with the ATP assay), HepG2 cells, and PTELC, with an IC_50_ of approximately 15 µM. This value aligns with data reported for several other cell lines, as summarized by Cwiklińska-Jurkowska et al. [61]. In contrast, PCKS and NRK cells showed moderately higher IC_50_ values around 35 µM, consistent with findings from other laboratories using rat PCKS [62] and the human kidney cell line HK-2 [63]. The IC_50_ values of over 100 µM that we observed in our PCKS experiments using the MTT assay are also reported in literature [64]. Importantly, our most sensitive models appear to be predictive of *in vivo* nephrotoxicity, as cisplatin plasma concentrations ranging from 7 to 20 µM in cancer patients have been associated with kidney injury [65, 66], and in mice, a plasma level of approximately 2 µM has been shown to significantly increase blood urea nitrogen (BUN), a marker of kidney injury [67].

Arsenic trioxide induced comparable IC_50_ values around 6 µM in PCKS and PCLS, and kidney cells, aligning with previous data in kidney cells [68, 69]. Embryonic kidney cells (HEK293) exhibited a ten times higher sensitivity with an IC_50_ of 0.5 µM which coincides with the peak plasma concentration measured in patients treated with arsenic trioxide for acute promyelocytic leukemia [70, 71]. In contrast, HepG2 cells were less sensitive (IC_50_ around 100 µM) consistent with literature data [72, 73], additionally highlighting cell line-specific differences in arsenic trioxide handling.

Aristolochic acid I elicited the most pronounced tissue-specific difference. PCLS were highly sensitive (IC_50_ around 5 µM), whereas PCKS exhibited significantly lower sensitivity within the tested concentration range. This is consistent with the hepatic predominance of enzymes responsible for bioactivating aristolochic acid I to its nephrotoxic metabolite aristolactam I [74]. The differential metabolic capacity between PCLS and PCKS likely explains this observation.

Among the potential negative control substances, melatonin and acarbose showed no cytotoxicity in MTT assays but elicited ATP reduction. This phenomenon, also reported in long-term studies on melatonin [75, 76], may reflect metabolic modulation rather than outright cytotoxicity. Acarbose-induced tissue shrinkage parallels the hyperosmolar stress observed with supraphysiological glucose, suggesting an osmotic rather than toxic effect. Surprisingly, dexamethasone decreased MTT-based viability, aligning with reports of glucocorticoid-induced growth inhibition in certain cancer cell lines [77, 78].

A key strength of this study is the high reproducibility of viability assays across different experimenters over extended periods, demonstrated with acetaminophen as test compound. The concordance between ATP and MTT measurements for acetaminophen supports the robustness of our optimized protocol. While this consistency may not generalize to all compounds, the reproducibility achieved underscores a critical advantage of tissue slice models over traditional cell lines.

## Conclusion

We successfully simplified the culture and toxicological testing of PCKS and PCLS, eliminating the need for specialized hyperoxic equipment and employing near-physiological culture conditions. This makes the approach feasible for widespread adoption in standard laboratory settings. Our data demonstrate that PCLS are highly sensitive and predictive of hepatotoxicity, with IC_50_ values consistent with clinically relevant concentrations and animal models. PCKS showed comparable sensitivity to kidney cell lines and could potentially be further optimized - particularly by reducing slice thickness to enhance sensitivity.

In summary, PCTS represent a reliable, reproducible, and physiologically relevant platform for nephrotoxicity and hepatotoxicity testing. Beyond improving concentration estimates for *in vivo* studies, this approach supports refinement and reduction principles in animal experimentation, contributing to more ethical and cost-effective research and test pipelines.

## Author contributions

N.S. designed the study, C.S., G.B., A.E., S.He., A.D., Z.R., F.R. and I.M.M.-J. performed research and analyzed the data, N.S., I.M.M.-J., G.F. and S.Ha. drafted and revised the manuscript, all authors approved the final version of the manuscript.

## Acknowledgment

The outstanding technical assistance of Kerstin De Mezzo is acknowledged.

